# Association between electrophysiological phenotypes and Kv2.1 potassium channel expression explained by geometrical analysis

**DOI:** 10.1101/2023.12.20.572720

**Authors:** Julio César Reyes-Garibaldi, Marco Arieli Herrera-Valdez

## Abstract

Excitable cells exhibit different electrophysiological profiles while responding to current stimulation in current-clamp experiments. In theory, the differences could be explained by changes in the expression of proteins mediating transmembrane ion transport. Experimental verification by performing systematic, controlled variations in the expression of proteins of the same type (e.g. voltage-dependent, noninactivating Kv2.1 channels) is difficult to achieve in the absence of other changes. However, biophysical models enable this possibility and allows us to assess and characterise the electrophysiological phenotypes associated to different levels of expression of non-inactivating voltage-dependent K-channels of type Kv2.1. To do so, we use a 2-dimensional biophysical model of neuronal membrane potential and study the phase plane geometry and bifurcation structures associated with different levels of Kv2.1 expression with the input current as bifurcation parameter. We find that increasing the expression of Kv2.1 channels reduces the size of the region of the phase plane from which action potentials can be initiated. The changes in expression can also be related to different transitions between rest and repetitive firing in current clamp experiments. For instance, increasing the number of Kv2.1 channels shifts the rheobase current to higher levels, but also expands the dynamic range in which excitatory external current produces repetitive spiking. Our analysis shows that changes in the responses to increasing input currents can be associated to different sequences of fixed point bifurcations. In general, the fixed points are attracting, then repulsive, and later become attracting again as the input current increases, but the bifurcation sequences also include changes in fixed point type, and change qualitatively with the expression of Kv2.1 channels. In the non-repetitive spiking regime with low current stimulation, low expression of Kv2.1 channels yields bifurcation sequences that include transitions between 3 and 1 fixed points, and repetitive firing starts with delays that decrease with increasing current (aggregation). For higher expression of Kv2.1 channels there is only one fixed point that changes in type and attractivity as the input current increases, convergence to rest tends to be oscillatory (resonance), and repetitive spiking starts without noticeable delays. Our models explain how the same neuron is theoretically be capable of including both aggregating and resonant modes of integration for synaptic input, as shown in current clamp experiments.

## 1 Introduction

Electrical signalling is important for fast communication between cells in all living organisms (Joukar, 2021; Sibaoka, 1991). Of particular interest, action potentials (APs) are electrical pulses of the transmembrane potential, or membrane potential for short, with amplitudes of tens to hundreds of millivolts (mV). The animal cell types known to produce APs include invertebrate and vertebrate neurons (Bean, 2007), glia (Fields, 2008), cardiac and striated muscles (Bean and Rios, 1989; Juel, 1988), and vertebrate pancreatic *β*-cells (Mattews and Sakamoto, 1975).

APs are produced by a nonlinear combination of electrodiffusive (also nonlinear) transmembrane currents carried by ions, which are mediated mainly by channel proteins (Herrera-Valdez and Lega, 2011); pumps also contribute to change the membrane potential, but their peak currents are at least one order of magnitude smaller in comparison to the peaks of the large currents (Benarroch, 2011; Gadsby, 2009). If we regard electrical excitability from here on as the capability of cells to produce APs, then we shall call a cell excited while it produces an AP, and sustained firing of APs can be thought of as a continued excited state. Since ion channel expression affects the way in which APs are created, it affects excitability. For a start, we focus on neuronal excitability and how it changes with the expression of K^+^channels, ubiquitous in the nervous system and other excitable tissues. There are different types of K^+^channels, with different biophysical properties that exert different effects on the membrane potential. In this paper we study how non-inactivating, voltage-dependent K^+^channels similar in function to Shab channels in Drosophila (Tsunoda and Salkoff, 1995) or the homologous subfamily of Kv2.1 channels in mammals (Murakoshi and Trimmer, 1999). We choose these types of channels because the belong to the most diverse family of voltage-gated ion channels studied up to date (Choe, 2002), and because they are mostly conserved across different excitable cell types in different invertebrate and vertebrate species (Salkoff et al., 1992).

At present, it is nearly impossible to perform systematic variations in the expression of a specific group of proteins of the same type without changing the expression of any other protein. For instance, variations in the expression of any subset of proteins may result in different, unpredictable numbers of other proteins, but may also yield proteins with possibly very different biophysical properties in comparison with wild types. One example of the latter is the heterogeneities in half-activating potentials from splice variants of voltage-gated sodium channels expressed from the same precursor genes (Lin et al., 2009). Nevertheless, biophysical models enable the possibility of studying the changes in neuronal excitability induced by changing the expression of Kv2.1 channels without any further genetic changes.

Experimental evidence shows that populations of neurons of the same type exhibit different degrees individual heterogeneity that can be quantified by examination of their excitability profiles. These profiles are typically defined by comparing current clamp electrophysiological recordings (Mason et al., 2005). Among other factors, the differences are believed to occur because of variations in the expression of proteins mediating ion transport across the membrane (Herrera-Valdez et al., 2013). Indeed, the way that neurons change between being excitable to a continuously excited state may vary in different ways depending on the levels of expression of different types of K-channels (Peng and Wu, 2007), especially Kv2.1 (Misonou et al., 2005).

The neuronal excitability resulting from different patterns of K^+^ channel expression that includes Shab (non-inactivating) and Shaker (inactivating) channels has been previously characterised from a statistical stand point using experimental data obtained from recordings in Drosophila (Peng and Wu, 2007) and in rats among other species (Mohapatra et al., 2009). However, it is important to distinguish different excitability profiles from a non-statistical perspective, via bifurcation analysis. Among other reasons, the dynamical systems approach allows the possibility of linking patterns of channel expression directly to the biophysics and geometry underlying the dynamics of the membrane potential, especially at transitions between different behaviours (bifurcations) displayed by the same neuron in response to current stimulation. There are also theoretical studies that aim to characterise neuronal excitability in terms of general mechanisms with dynamical systems approaches based on bifurcation analysis using conductance-based models (Av-Ron et al., 1991; Av-Ron et al., 1993; Izhikevich, 2007; Rinzel and Ermentrout, 1989). A subset of such studies has focused on analysing excitability varying the contribution of inactivating, voltage-gated A-type K^+^ channels (Av-Ron, 1994; Drion et al., 2015).

We start addressing how does the expression of K^+^ channels change the excitability of a neuron. To do so, we use a basic 2D biophysical model with a minimum complement of three voltage-dependent currents mediated by Na-K ATPases, transiently activated Na channels, and Kv2.1-like channels. This is an important and necessary point of departure before considering questions involving more complicated membrane profiles of channel expression (e.g. Drion et al., 2015) that would imply higher dimensionality and more parameters. The model used here was derived from biophysical principles without making assumptions about electrical circuits (Herrera-Valdez, 2014, 2018, 2020) and improves over a number of issues in other biophysical models (e.g. conductance-based, see Herrera-Valdez (2012a) and discussion). The general idea is to perform a systematic variation in the expression of non-inactivating voltage-dependent K-channels, and analyse the behaviour of the model in current clamp situations while leaving other parameters fixed. One of our objectives is to explain in more detail the changes that can be observed from the same model neuron in response to current clamps of different, constant amplitudes, for different expression profiles of Shab-like or Kv2.1-like channels as reported in the literature (Peng and Wu, 2007). In particular, we are interested in describing mechanisms by which the same neuron can integrate inputs in resonant and non-resonant ways (Destexhe et al., 2003; Izhikevich, 2007). For instance, in current clamp experiments it can be observed that the *same* neuron may display repetitive spiking after a delay for some level of external current stimulation, stop showing the delay for larger amplitudes of the same kind of stimulus, and show depolarisation block for even larger stimuli (Granit et al., 1963a,b; Peng and Wu, 2007). How would these transitions occur for different profiles of Kv2.1 channel expression?

### 1.1 Membrane potential and electrical excitability

The ion fluxes that change the membrane potential can be thought as two types: those mediated by pumps that transport at least one type of ion across the membrane against its electrochemical gradient (Blaustein et al., 2004; Gadsby, 2009), and those mediated by ion-selective channels that mediate passive, electrodiffusive transmembrane fluxes (Hille, 1992). Ion channels are gated by different factors that include voltage (Bezanilla, 2007), ligands (Collingridge et al., 2009) such as Ca^2+^ (Petersen and Maruyama, 1984; Sah, 1996), Na^+^(Dryer, 1994), and glutamate (Wo and Oswald, 1995); or possibly other factors like light (Shi et al., 2019). Pumps mechanically translocate ions across the membrane at rates between 3 and 7 orders of magnitude slower than the electrodiffusive transport rates for channels (Gadsby, 2009; Stein and Litman, 2014). Nonetheless, the electrogenic activity of pumps is one of the main determinants of the resting membrane potential and the overall excitability in neurons (Wang et al., 2012; Wright, 2004). Of note, the Na-K ATPase is arguably the most important ionic pump in nature. It is the main driver of primary active transport of Na^+^ in cells, moving 3 Na^+^ions outward in exchange for 2 K^+^ions moving inward, both against their electrochemical gradient (Munakata et al., 1998). The energy for the transport is obtained from the hydrolisation of ATP, which causes the phosphorylation of an aspartate residue that is highly conserved across different cell types in different species (Skou and Esmann, 1992), and enables a conformational change that mediates the electromechanical exchange.

Neuronal APs are upward pulses in the membrane potential starting with an upstroke that typically reaches a maximum depolarisation rate of 100 mV/ms or more, with a slightly slower downstroke (Naundorf et al., 2006; Villiere and McLachlan, 1996). The upstroke is mainly produced by large inward transmembrane negative currents typically carried by Na^+^(Stuart and Häusser, 1994), Ca^2+^(Zang and De Schutter, 2021), or both (Bean, 2007), depending on factors that include the neuronal compartment producing the AP and the cell type. The downstroke is mainly produced by positive outward currents typically carried by K^+^(Murakoshi and Trimmer, 1999; Speca et al., 2014; Tsunoda and Salkoff, 1995), and possibly negative inward currents carried by Cl^−1^. Nevertheless, the combination and timing of the ion fluxes that cause an AP may vary depending on the cell type and the organism.

### 1.2 Experimental current clamps and families of dynamical systems

#### Current clamp (I-clamp)

Experiments allow us to sample the diversity of behaviours that can be displayed by a single neuron (Herrera-Valdez et al., 2013; Hounsgaard et al., 1984). I-clamp experiments consist in recording the transmembrane potential of a cell in response to current stimulation. One of the most common current clamp protocols consists of injecting square-shaped current pulses in three phases in which the stimulus amplitude is zero during the first and third phase, and the amplitude is set to a different constant value during the second phase. A whole recording typically consists of one or more trials, each involving a sequence *I*_0_ < … < *I_n_* of current amplitudes injected during the second phase of the different trials. Excitable cells (e.g. neurons) that do not exhibit spontaneous sustained firing in the absence of a stimulus (e.g. zero current in I-clamp) have a resting level that may vary between −85 and −50 mV approximately depending on the cell type (Aidley, 1998; Suter et al., 2013). The amplitude of the stimulus during the second phase of the I-clamp can be such that the membrane converges to a resting voltage, but such a voltage changes according to the stimulus in comparison to the resting potential for zero current. If that is the case, the voltage to which the membrane converges can be thought of as one of the coordinates of an attracting fixed point in a dynamical system. Alternatively, the amplitude of the stimulus may be such that the membrane potential displays repetitive firing. From a dynamical systems perspective, this is a different kind of attractor called limit cycle.

#### Neuronal membrane dynamics

The neuronal membrane potential can be modelled with a non-linear dynamical system with a minimum of two variables, say *v* and *w*, respectively representing the membrane potential and the proportion of open delayed rectifier (non-inactivating or slowly inactivating) channels (Av-Ron et al., 1991; Rinzel, 1985; Rinzel and Ermentrout, 1989; Tsunoda and Salkoff, 1995). For APs to occur, there must be a separation of time scales for the two variables of the system (Ermentrout and Terman, 2010; FitzHugh, 1961), with *v* displaying self-amplifying and faster dynamics in comparison to *w*. In contrast, *w* should provide negative feedback to *v* with slower dynamics (Izhikevich, 2007). Explicitly, these properties can be formulated by first setting a nonlinear, autonomous system of ordinary differential equations the form

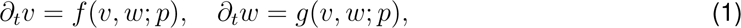

where *∂_t_v* represents the instantaneous rate of change with respect to time, *f* and *g* are nonlinear functions, and *p* is a vector of parameters.

Assuming that the biophysical properties of the recorded neuron do not change during an I-clamp experiment (e.g. the ion channels of any given type do not change in number), the current amplitudes can be thought of as different values for one of the entries in the parameter vector *p*. Therefore, to capture the variability in the voltage responses for different current amplitudes, it is necessary to model the behaviour of neuron with not one, but a *family* of autonomous dynamical systems

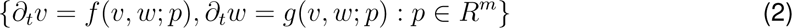

with the same functional forms for *f* and *g*, and different values of the stimulus current, in the absence of other changes. In such a family, there would be components in the evolution rules *f* and *g* that are common to all members of the family and represent the intrinsic, *core* components for the dynamics of a single neuron. The non-common components can be thought of as representing the experimental variations that give rise to qualitatively different behaviours produced by the same neuron. If on top of the experimental manipulations we wish to take into account populations of neurons, then *p* could also include differences in the biophysical properties for different neurons. More specifically, one or more of the entries of *p* could represent different expression patterns for ion channels (Herrera-Valdez et al., 2013; Misonou et al., 2005).

## 2 Materials & Methods

### 2.1 Modelling

#### Dynamics

Consider a system modelling a patch of plasma membrane and its surrounding extra and intracellular bulk compartments. Assume that he intra and extracellular compartments are isopotential and that the total charge density of the system is constant with respect to time. That is, 0 = *∂_t_* (*Q_A_* + *Q_M_*), where *Q_A_* and *Q_M_* represent the charge densities around, and crossing through the membrane at any given time. Assume further that the accumulation of charge at the membrane-liquid interface is a monotonically increasing function of *v*, the transmembrane potential. As a consequence,

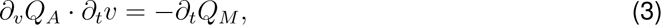

with *∂_t_Q_M_*representing the transmembrane current density (in pA/*µ*m^2^, time in milliseconds), which can be written as a sum of an external current *I_F_* and transmembrane fluxes of Na^+^and K^+^, with current densities *I*_NK_, *I*_KD_, *I*_NT_, mediated respectively by Na-K ATPases, voltage-dependent K^+^delayed rectifier channels, and transiently activating voltage-dependent Na^+^channels, respectively.

Notice that the dynamics in equation (3) are not derived by assuming any electrical circuit analogies (Herrera-Valdez, 2020). However, assuming that *Q_A_* = *Cv* with constant *C* (in pF/*µ*m^2^), yields the classical formulation derived from an “equivalent” electrical circuit (Hodgkin and Huxley, 1952). In this case *C* can be thought of, as the membrane “capacitance” from the equivalent circuit model, and equation (3) can be rewritten as

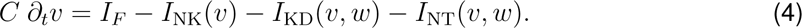

with *v ∈* [*v*_KD_*, v*_NT_] [*−*90, 60] mV, and *w ∈* [0, 1]. Varying *I_F_* within a reasonable interval that resembles the range for stimulus amplitudes from I-clamp experiments defines at least one family of dynamical systems representing a single cell in the absence of changes. That is, the parameter vector 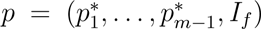 could be varied in only the dimension of *I_F_* to reproduce the responses of a single neuron in I-clamp experiments.

It will be assumed that all the K-channels in a single neuron model have the same biophysical properties. Nevertheless, variations in the number of potassium channels expand the family in a second (co)dimension and could be interpreted as changing the neuron, as such changes would take too long in comparison to the duration of a voltage-clamp experiment. Therefore, the family of dynamical systems obtained by varying *I_F_*, or any that does not change the biophysical properties of the model can be regarded as a single neuron.

Each of the transmembrane currents mediated by ions can be written using a general formulation for transmembrane transport derived from thermodynamical principles (Herrera-Valdez, 2018). The thermodynamic transmembrane transport model makes use of two exponential functions of the form

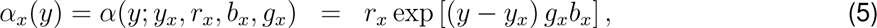

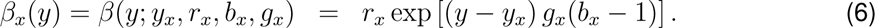

The functions *α_x_* and *β_x_* represent forward and backward fluxes mediated by membrane proteins of type *x* (e.g. Na-K ATPase), or alternatively, forward and backward rates for the conformational change responsible for the gating of a channel (e.g. *w* representing K-channel activation or *m* representing NaT-channel activation). Therefore, the label *x* represents any of the labels *{*KD, NT, NK*, m, w}*. Of note, *b_x_* captures the bidirectional, but not necessarily symmetrical nature of the transmembrane fluxes that underlies the rectification (Hodgkin and Horowicz, 1959; Nakajima et al., 1961, 1962), and similarly, possibly asymmetrical gating rates. More specifically, for ion fluxes, *r_x_*, *b_x_*, and *y_x_* respectively represent the single-protein current amplitude (pA/*µ*m^2^), *b_x_* (in [0, 1]) the bidirectional but not necessarily symmetrical nature of the transmembrane fluxes that underlies rectification (Hodgkin and Horowicz, 1959; Nakajima et al., 1961, 1962), and the reversal potential for the flux (mV). For channel gating, *r_x_* controls the rate of the reaction underlying the gating conformational change (1/ms), *b_x_* the bias in the forward direction of the reaction (in [0, 1]), *g_x_* is a gain control for the gating, and a reversal electromechanical potential (see Table 1 for parameter values). The potentials in the model would be then be normalised by the thermal potential *v_T_* = *kT/q* (in mV)^1^.

**Table 1:** Model parameters. Details on the derivation of the transport formulas can be found in the work by Herrera-Valdez (2018).

| Parameter | Value | Units | Description |
| --- | --- | --- | --- |
| $a_{KD}$ | [4,20] | $\mu\text{m}^2/(\text{mV pF})$ | K-channel population flux rate |
| $a_{NT}$ | 4 | $\mu\text{m}^2/(\text{mV pF})$ | Na-channel population flux rate |
| $a_{NK}$ | 800 | $\mu\text{m}^2/(\text{mV pF})$ | Na-K ATPase population flux rate |
| $r_{KD}$ | 1 | $\text{pA}/\mu\text{m}^2$ | Single K-channel transport rate |
| $r_{NT}$ | 1 | $\text{pA}/\mu\text{m}^2$ | Single Na-channel transport rate |
| $r_{NK}$ | $10^{-4}$ | $\text{pA}/\mu\text{m}^2$ | Single Na-K ATPase transport rate |
| $r_w$ | 0.25 | $\text{ms}^{-1}$ | K-channel gating activation rate |
| $l$ | 0.35 | – | K-channel cooperativity (logistic-like dynamics) |
| $g_w$ | 2.1 | – | K-channel activation gain control |
| $g_m$ | 4 | – | Na-channel activation gain control |
| $b_w$ | 0.65 | – | K-channel activation forward bias |
| $b_{NT}$ | 0.5 | – | Na-channel forward flux (inward) bias |
| $b_{KD}$ | 0.3 | – | K-channel forward flux (outward) bias |
| $b_{NK}$ | 0.1 | – | Na-K ATPase forward bias |
| $v_w$ | -10 | mV | K-channel half-activating potential |
| $v_m$ | -15 | mV | Na-channel half-activating potential |
| $v_{NT}$ | 60 | mV | $\text{Na}^+$ Nernst-potential |
| $v_{KD}$ | -90 | mV | $\text{K}^+$ Nernst-potential |
| $v_{NK}$ | -80 | mV | Na-K ATPase reversal potential |
| $v_T$ | 26.64 | mV | Thermal potential |
| $C$ | 20 | $\text{pF}/\mu\text{m}^2$ | Membrane "capacitance" |

The general functional form for the currents in the thermodynamic model is

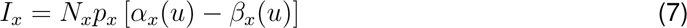

where *x* is a label representing the transmembrane transport mechanism of interest, with *N_x_* representing the number of transmembrane proteins mediating the flux and *p_x_* in [0,1] representing the proportion of active membrane proteins of type *x*. All transmembrane concentrations of ions and the ATP/ADP ratio are assumed to be constant (see Johnston et al., 1995, chapter 2, example 2.1). As a consequence, the Nernst potentials *v*_Ka_ and *v*_Na_ for K^+^and Na^+^, the hydrolisation potential *v*_ATP_, and the reversal potential *v*_NK_ = *v*_ATP_ + 3*v*_Na_ *−* 2*v*_Ka_ of the Na-K ATPase are all constant (for details of the biophysical derivation from first principles and relevant literature see Herrera-Valdez, 2018). As already mentioned, the absolute amplitude *I*_NK_ is smaller by at least one order of magnitude with respect to the absolute amplitudes for *I*_KD_ and *I*_NT_.

The activation of K^+^channels will be represented by a variable *w* with dynamics given by

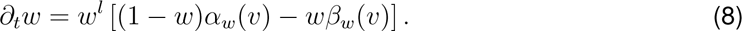

It is assumed that *w* also represents the inactivation of Na^+^channels. As a consequence, 1 *− w* represents the proportion of Na^+^channels that are not inactivated.

#### Changes of variables and reparametrizations

To simplify the notation and computational expense, we work with the variable *u* = *v/v_T_* and the parameters *u_x_* = *v_x_/v_T_*, for *x ∈* {NK, KD, NT*, w, m*}. After re-scaling *v → u* = *v/v_T_* and dividing by *C*, the evolution for the membrane dynamics can be rewritten as

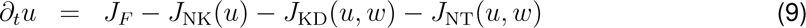

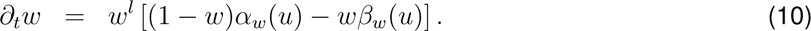

As a consequence, each of the *J_x_* is in units of (1/ms), and the re-scaled currents given by

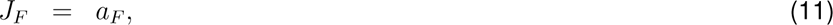

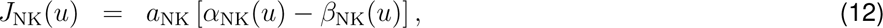

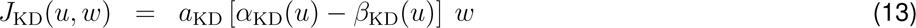

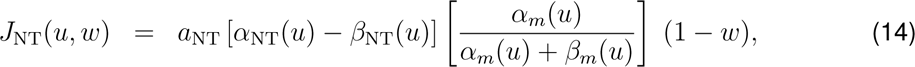

and grouped parameters *a_F_* = *I_F_ /*(*v_T_ C*) and *a_x_* = *N_x_/*(*v_T_ C*) in (mV pF/*µ*m^2^)^−1^, for *x ∈* (NK, KD, NT). The last terms in equations (13)-(14) correspond to *p*_KD_(*v*) and *p*_NT_(*v*), the voltage-dependent gating for the K^+^and Na^+^channels, and *p*_NK_ = 1 (see equation (7)).

The idea for this article is to exploit the biophysical formulation re-scaled in equations (9)-(14) to study the geometry underlying the transitions between rest, repetitive firing, and depolarisation block in current-clamp stimulation protocols taking into account the underlying sequences of bifurcations associated to changes in the expression of Kv2.1 channels.

#### Notes on parameter choices, experimental values, and illustrations

The construction hypotheses and parameter regimes for the neuronal dynamics in this paper follow criteria that include

- Kv2.1 channel activation. The time course of *w* in a voltage-clamp should be sigmoidal, akin to solutions of logistic equations (see Fig. 3 in the seminal work of Hodgkin and Huxley (1952) and other researchers including Covarrubias et al. (1991); Tsunoda and Salkoff (1995)), with convergence of *w* toward its steady state within 10-15 milliseconds, which means that the time constant for *w* should be approximately between 3-5 ms. This requirement can be satisfied by assuming evolution rules for *w* that resemble logistic dynamics, in agreement with the idea that *w* should be modelling gating at the population level.
- Na-K ATPases. The resting potential depends on the reversal potential for Na-K ATPases (Ashcroft, 2005). The rate of transport of Na-K ATPases should be between 3 and 4 orders of magnitude (Heyse et al., 1994) smaller in comparison to the large K^+^and Na^+^currents mediated by channels in the model. The reversal potential for Na-K ATPases currents mediated should be between −90 and −60 mV (Chapman, 1978; Skou and Esmann, 1992), which in turn depends on ion-concentration gradients (Chapman, 1973) and the voltage associated to ATP-hydrolysis (Herrera-Valdez, 2018). These requirements can be met by selecting appropriate values for *r*_NK_, and *v*_NK_ with *a_F_* = 0 and of crucial importance to obtain realistic dynamics around rest.
- The maximum *∂_t_v* should be at least 100 V/s. This requirement should be met by setting large enough values for *a*_NT_ (or equivalently, *N*_NT_). Also, the downstroke should be at least half as fast as the upstroke (2*|* min *∂_t_v| < |* max *∂_t_v|*) (Carter and Bean, 2009; Suter et al., 2013, among other researchers), which involves setting *b*_KD_, the rectification bias for K-channels to values lower than 1/2 to yield inward rectification.
- The duration of action potentials for this model should be between 0.5 and 3 milliseconds (Carter and Bean, 2009). This requirement can be met by adjusting *r_w_*, mainly.
- The rheobase, understood here as the minimum amplitude for sustained current stimulation required to observe repetitive spiking, should be at least 50 pA (varying *I_F_*). To determine the rheobase for different instances of our model we will restrict our calculations to 1 second stimuli and use a maximum resolution of 1 pA.

#### Numerics

Numerical calculations are performed in terms of *u* and *w*, but all illustrations will show the membrane potential *v* (instead of *u*); the idea is to keep the relevant biophysical variables in mind and interpret the results accordingly. To be able to draw comparisons with time courses of different trajectories, we will always show the membrane potential in the vertical axis. Numerical solutions were calculated using time steps between 1/40 and 1/70 ms. Computational models were implemented in Python 3.11(www.python.org) with modules sympy (http://www.sympy.org), numpy (http://numpy.org), and matplotlib (http://www.matplotlib.org), using personal computers.

## 3 Results

From here on each model neuron will be defined as a family of dynamical systems in which the input current varies, while leaving all other parameters the same (i.e. biophysical properties are kept constant for each neuron). By extension, a population of model neurons displaying different levels of expression for Kv2.1 channels would be an even larger family in which the parameter *a*_KD_ varies too. Recall that we regard a single dynamical system as excitable if it has the capability of displaying action potentials from some subset of initial conditions.

**One box-shaped current clamp, three dynamical systems.** Notice that if we subject a model neuron to current stimulation as would be the case in a current-clamp experiment with a box-shaped protocol, during the first phase of stimulation with zero current the state of the system should approach the closest attractor to the initial condition. During the second phase of stimulation the evolution of the change of the stimulus current amplitude changes the system to a new autonomous regime, the (prior) near-steady state of the system becomes an initial condition, and the system evolves from there to the closest (new) attractor. Excitation takes place if the response to the stimulus includes action potentials. The system would go back to its original dynamics for the third phase, evolving from whatever state was visited prior to the return to zero current stimulation. The system would then evolve according to the original rules for zero current, but from an initial condition defined by the last state visited prior to the return to zero current.

We now focus on geometrical properties that explain how excitability is shaped by the expression of Kv2.1 channels and then use the box-shaped current clamp protocol with zero current for start and end phases as a template for stimulation and analysis.

### 3.1 Association between the expression of potassium channels and the geometry and dynamics of the model

For an initial analysis, let us consider the phase plane, vector field, and nullclines for a system given by equations (9)-(14) for (*w, u*) *∈ D* = [0, 1] *×* [*u*_K_*, u*_Na_].

The *w-nullcline* is the set states *w*_Null_ = {(*w, u*) : *∂_t_w* = 0}, that includes the graph of the function

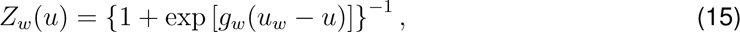

(Fig. 1, orange curves, after re-scaling *v* = *uv_T_*) with the line {(0*, u*) : *u ∈* [*u*_K_*, u*_Na_]} if *l >* 0 (not shown in Fig. 1). Notice that the *w*-nullcline is not affected by solely changing the number of K-channels in the model.

**Figure 1:**
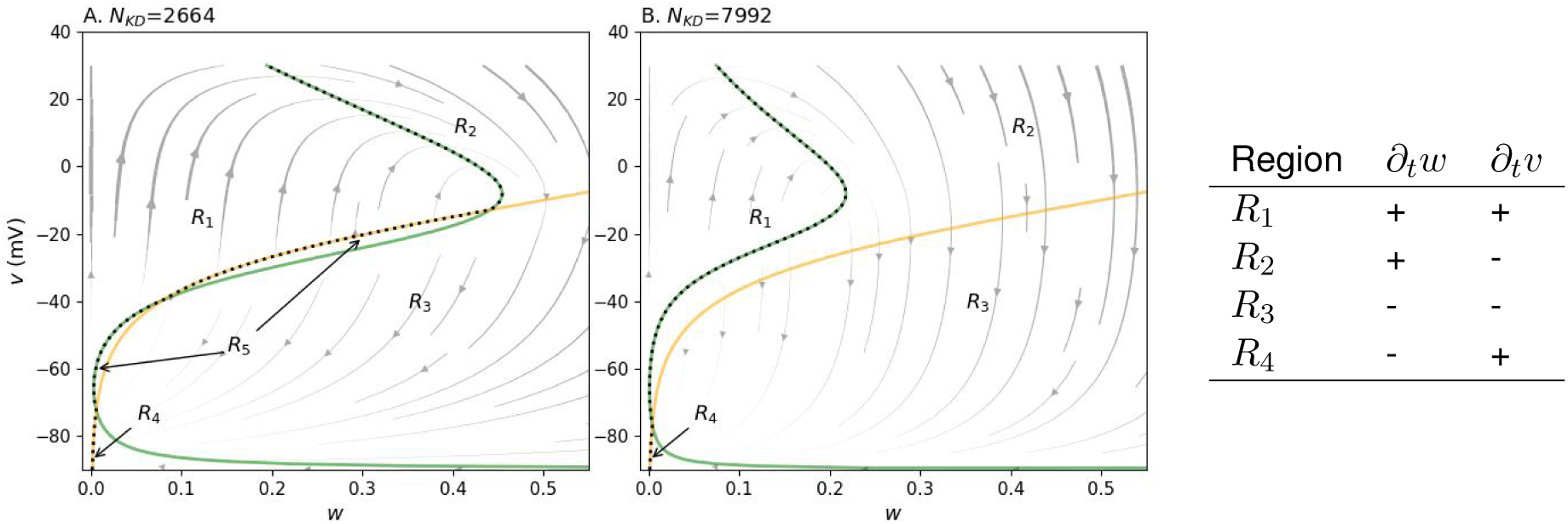
Phase planes and local dynamics. A and B Phase plane corresponding to *N*_KD_ = 2664 and *N*_KD_ = 7992, respectively. The green and orange curves are the *v*- and *w*-nullclines, respectively. The grey lines with arrows represent trajectories. The different regions defined in the table are shown for each case. The curve in black dots shows the upper boundary in the *w*-direction for region *R*_1_.

The *u*-nullcline *u*_Null_ = {(*w, u*) : *∂_t_u* = 0} can also be thought of as the graph of a function

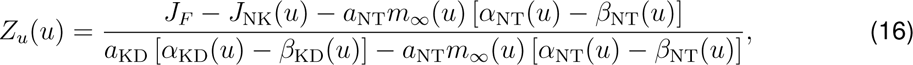

whose graph has the shape of a cubic function (Fig. 1, green curve. Also see FitzHugh, 1961).

The function *Z_u_*(*u*) has three branches which will be referred to as lower, middle, and upper branches, separated by a local minimum and a local maximum, respectively. The upper and lower branches are such that *∂_u_Z_u_ <* 0, and in the middle branch *∂_u_Z_u_ >* 0. Notice that increasing the input current *J_F_* increases the *w*-values of *Z_u_*, thereby affecting the location and possibly the type of the fixed points in the set {(*w_∗_, u_∗_*) : *∂_t_u* = 0 = *∂_t_w*}. In addition, increasing the number of Kv2.1 channels in the model (*a*_KD_ in the denominator in equation (16)) decreases the effects caused by an increase in *J_F_*. In other words, increasing the input current will translate into a right shift in the *w*-direction for the *u*-nullcline of size *a_F_* (*a*_KD_ [*α*_KD_(*u*) *− β*_KD_(*u*)] *− a*_NT_*m_∞_*(*u*) [*α*_NT_(*u*) *− β*_NT_(*u*)])^−1^, which decreases as *a*_KD_ increases, and becomes more important for values of *u* larger than rest. As a consequence, increasing *a*_KD_ in the (model) neuron would mean to decrease the overall impact of the input current, and in particular, increase the rheobase.

Within the relevant biophysical ranges for the parameters in the model, the number of fixed points in systems like these can be 1, 2, or 3 (Fig. 1, A-B). The (rare) case in which there are two fixed points only occurs at the transition between one and three fixed points. One way to observe the transition is to increase the number of Kv2.1 channels in the model, which decreases the all *w*-values of *u_Null_*, thus lowering the local maximum of *Z_u_* near *u* = 0. The *u*-value at which *Z_u_* reaches a local *w*-minimum and the slope of the middle branch in *u_Null_* also decrease. Another way in which the transition from 3 to 1 fixed points can be observed is to increase the value of *g_w_*, which controls the slope of the *w*-nullcline.

#### Trajectories and excitability

The discussion below applies only to autonomous dynamical systems modelling excitable phenomena. Without loss of generality, the phase plane *w*-*v* can be divided into four regions labelled *R*_1_,…,*R*_4_ according to the signs of the components of the vector field at each point (Fig. 1), with *R*_1_ representing the region where *∂_t_w >* 0 and *∂_t_v >* 0, *R*_2_ such that *∂_t_w >* 0 *> ∂_t_v*, *R*_3_ where *∂_t_w <* 0 and *∂_t_v <* 0, and *R*_4_ such that *∂_t_w <* 0 *< ∂_t_v*. Notice that *R_i_*, with *i* = 1*,..,* 4 represent all the possible non-horizontal and non-vertical direction fields a phase plane can have (Rinzel and Ermentrout, 1998, see). If there were three fixed points, there would be a pair of extra regions delimited by the nullcline segments joining the three fixed points, with dynamics equivalent to those of *R*_2_ (*∂_t_v <* 0 *< ∂_t_w*) and *R*_4_ (*∂_t_w <* 0 *< ∂_t_v*), respectively (Table in Fig. 1, panel A).

Taking the observations above into account, the excitability of an autonomous system can be defined only in terms of the trajectories that it can produce from different initial conditions (Fig. 2). A list of necessary conditions to observe excitability in a 2-dimensional autonomous, continuous, dynamical system like the one from equations (9)-(14) should include that: (a) all of the four regions

**Figure 2:**
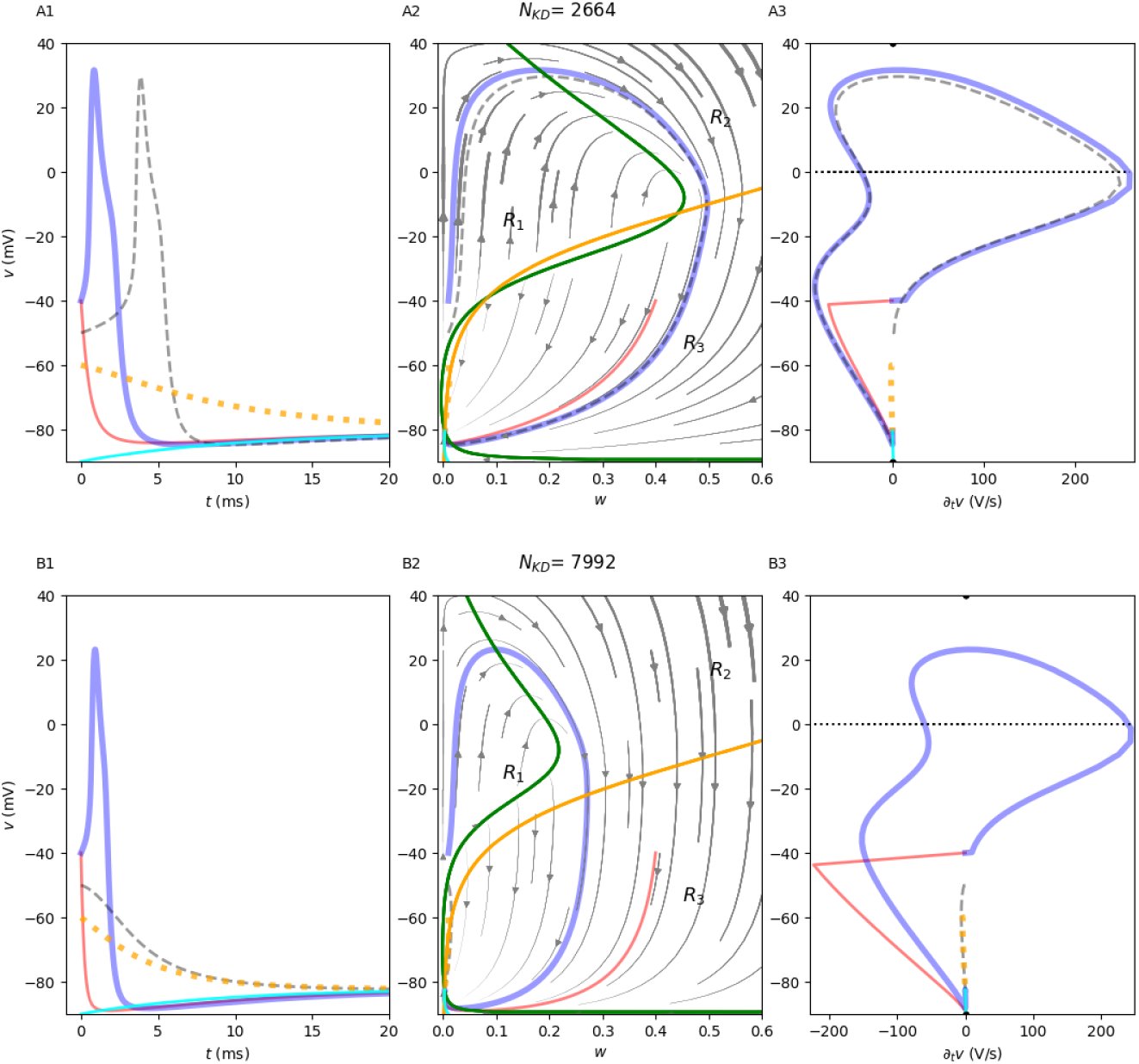
Dynamics of qualitatively different excitable systems displaying convergence toward attractor points for different levels of expression for K^+^channels (*N*_KD_ *∈ {*2464, 7992*}*, which yields 1 or 3 fixed points in A1-A3 and B1-B3, respectively). A1 and B1 show trajectories in the (*t, v*) plane. A2 and B2 show the phase planes with nullclines and the phase-plane trajectories that correspond to those shown in A1 and B1. The *v*- and *w*-nullclines are shown in green and orange lines, respectively. A3 and B3 show trajectories in the (*∂_t_v, v*) plane. All panels were obtained assuming zero input current, and the same set of initial conditions (*w*_0_*, v*_0_) (*v*_0_ *∈ {−*90*, −*60*, −*50*, −*40*}*, with *w*_0_ = 0.01 in blue, grey dashes, red dots, and cyan, respectively, (*w*_0_*, v*_0_) = (*−*40, 0.4) in red). All graphs are shown with *v* as a common vertical axis with the same range.

*R_i_*, *i ∈* {1*, …,* 4} exist, and surround the fixed point(s), located sequentially so that there are trajectories that can traverse *R*_1_, then *R*_2_, *R*_3_, and finally *R*_4_ in that order (or the reverse), before reaching an attractor. This means that any simple curve on the phase plane that encloses the fixed points and passes through *R*_1_, *R*_2_, *R*_3_, and *R*_4_, in that order, should have index −1 (or 1 if the order is inverted); (b) there is difference in characteristic time-scales of at least one order of magnitude in the region where both change in the same direction (say, *R*_1_, or *R*_4_ for down-going pulses); (c) the fast (“excitable”) variable *v* has self-amplifying properties, and the “slower” variable provides negative feedback to the fast variable; and last but most important. The separation of time scales should be such that *∂_t_v >* 10 *· ∂_t_w >* 0 somewhere in *R*_1_. We regard excitability as the capability of producing pulses, which for the case of neurons means producing action potentials. But what is a pulse? Let us adopt a (working) mathematical definition for an “upward” pulse: the fast variable of the system displays an upward pulse as a function of time, if the corresponding trajectory in the phase plane contains points in *R*_1_, that traverse into *R*_2_, then *R*_3_, and reach *R*_4_, in that order, at least once. An important distinction is in place here. The start of a pulse upward can occur without traversing the other three regions, as it happens with depolarisation blocks (Av-Ron et al., 1991; Rinzel, 1985). In such cases we will say pulsed are incomplete, and the system will not be regarded as excitable. Note that in such cases the system does have regions *R*_1_,…, *R*_4_ arranged as required before, but the system has at least a fixed point attractor located beyond the second branch of the *u*-nullcline. Pulses can thus back to rest, or occur repeatedly, as long as they are completed (traverse all four regions). Notice the that the upper bound in the *w*-direction for region *R*_1_ can be defined as

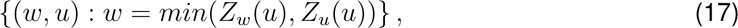

and used to identify initial conditions yielding action potentials. Similar conditions can be described for down-ward pulses (e.g. *∂_t_v <* 10 *· ∂_t_w <* 0).

As already mentioned, in these kinds of systems trajectories may either converge to a fixed point, or to a limit cycle. In fact, these systems may be bistable, often with one fixed point and one limit cycle as attractors. Should there be two attractor fixed points in a bistable system, the system would be unlikely to display pulses (see Süel et al., 2006, for an example of a non-neuronal excitable system). It might be possible that there are other combinations of attractors in systems like these (non of which we have knowledge of); the analysis in this study will not consider such cases.

Therefore, up to topological equivalence between phase planes and their vector fields, the regions *R*_1_,…,*R*_4_ in an excitable system like the one studied here are such that the trajectories starting in *R*_1_ will go through *R*_2_, then *R*_3_, and *R*_4_ before going to an attractor which may be a fixed point or a limit cycle (traversing the four regions periodically) (Av-Ron et al., 1991; Rinzel, 1985; Süel et al., 2006; Suel et al., 2007). For contrast, in the absence of a limit cycle attractor that traverses the regions *R*_1_, *R*_2_, *R*_3_, and then *R*_4_, those trajectories starting from regions that are not *R*_1_ do not result in APs (e.g. orange and cyan traces in Fig. 2A-C, but compare grey dashed traces in panel A with those of panels B and C). Therefore, we can regard *R*_1_ as the “excitability” region of the system, if the conditions (a)-(c) above are met. In particular, this means that trajectories starting in *R*_1_, go through *R*_2_, *R*_3_, and then *R*_4_ on their way to an attractor. In the absence of limit cycle attractors, initial conditions with high initial *v*-values do not necessarily yield APs. For instance, in the absence of a nearby limit cycle attractor, a trajectory may start at a large value for *v* outside of *R*_1_ and not display a pulse (e.g. orange traces in Fig. 2A-C).

#### Excitability for non-zero currents, bifurcations, and rheobase

Positive input currents shift the location of the *u*-nullcline toward higher *w*-values, thereby shifting the location of the fixed points to higher *w* values. For a family of dynamical systems modelling a single neuron, the rheobase can be thought of as a (codimension 1) bifurcation value of *I_F_* (equivalently, *a_F_*). Nevertheless, excitability is still a property that a single dynamical system may exhibit without requiring bifurcations.

#### Kv2.1 channels and steady state input current

The effects of increasing the number of Kv2.1 channels on the number of fixed points in the system can also be explored by examination of the steady state equation after solving for the input current. Explicitly,

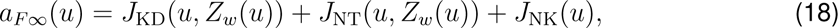

which results (Fig. 3 after re-scaling *I_F_* = *a_F_ v_T_ C*).

**Figure 3:**
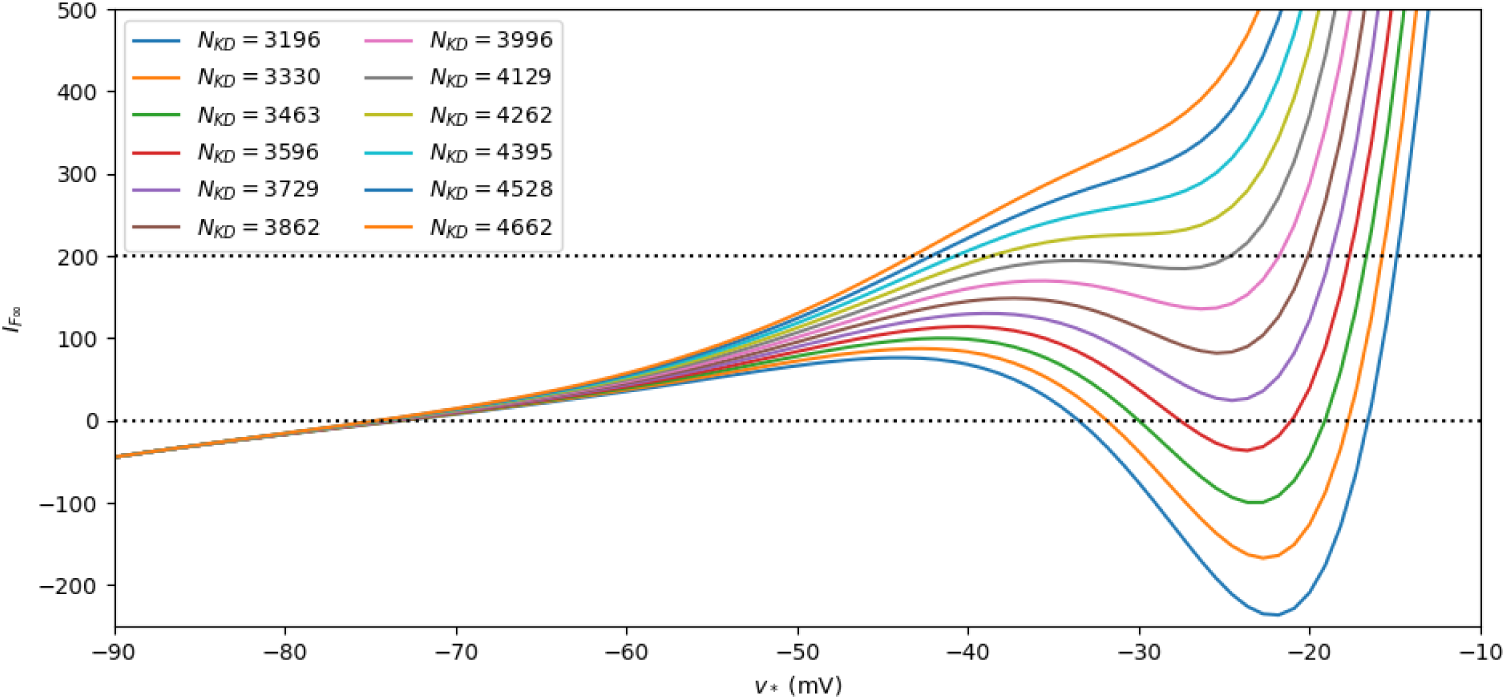
Saddle node bifurcation caused by the increase in *N*_KD_, illustrated by the steady state curve *a_F_*(shown after re-scaling *a_F_ → I_F_* = *a_F_ v_T_ C*. The horizontal black dotted lines illustrate two current amplitudes for which profiles of expression of Kv2.1 channels yield 3 fixed points.

It is not possible to analytically obtain *u_∗_*, the *u*-values from a fixed point, from *a_F_* using equation (18). However, it is possible to obtain values of *a_F_* (and therefore *I_F_*) from varying *u*. In other words, the pairs (*u_∗_, a_F∞_*(*u_∗_*)) satisfying the fixed point equation (18) yield the amplitudes of the (normalised) input current as a function of *u_∗_*. For instance, the roots of *a_F ∞_* (Fig. 3, black dotted horizontal line at *I_F∞_* = 0) correspond with the fixed points of the system at zero stimulation.

Also, it is possible to obtain bifurcation diagrams with *a_F_* as a bifurcation parameter without the need to numerically approximate the *u_∗_*-values. In particular, if *a_F∞_* is monotonically increasing as a function of *u*, then there is only one fixed point. In contrast, if *a_F ∞_* is not monotonic, then there are 3 fixed points.

Further, as the number of Kv2.1 channels increases (Fig. 3, different colour lines), the number of fixed points decreases from 3 to 1 and the transition occurs as two of the fixed points collapse and disappear; a typical saddle-node bifurcation.

### 3.2 Bifurcations taking Kv2.1 expression into account

We focus next on bifurcations associated to the expression of Kv2.1 channels and their influence on excitability. To do so, we analyse the fixed point bifurcation structures of families of dynamical systems defined by (9)-(14) with varying *a_F_* (codimension 1). Briefly, we set the expression of Kv2.1 channels to a particular level (while keeping the other biophysical properties constant), calculate *w_∗_* and *a_F_* (equations (15) and (18) respectively) from *u* within the range [*u*_KD_*, u*_NT_], determine the fixed point types and their attractivity (alt. repulsivity), and analyse the resulting fixed-point bifurcations to infer transitions in the local dynamics. This approach allows us to study the shape of the trajectories from different initial conditions. As before, illustrations show the original variables after re-scaling.

The steady state curves for different expression profiles of Kv2.1 channels (Fig. 4, transitions from filled to empty circles from left to right in each curve) show different sequences of bifurcations of fixed points. One common sub-sequence of bifurcations that can be observed as the input current increases, is that the fixed points with values around *v*_NK_ are attractors (associated to resting potentials and lower values of *I_F_*), then become repulsive (close to or at repetitive firing regimes), and then become attractors again (depolarisation blocks).

**Figure 4:**
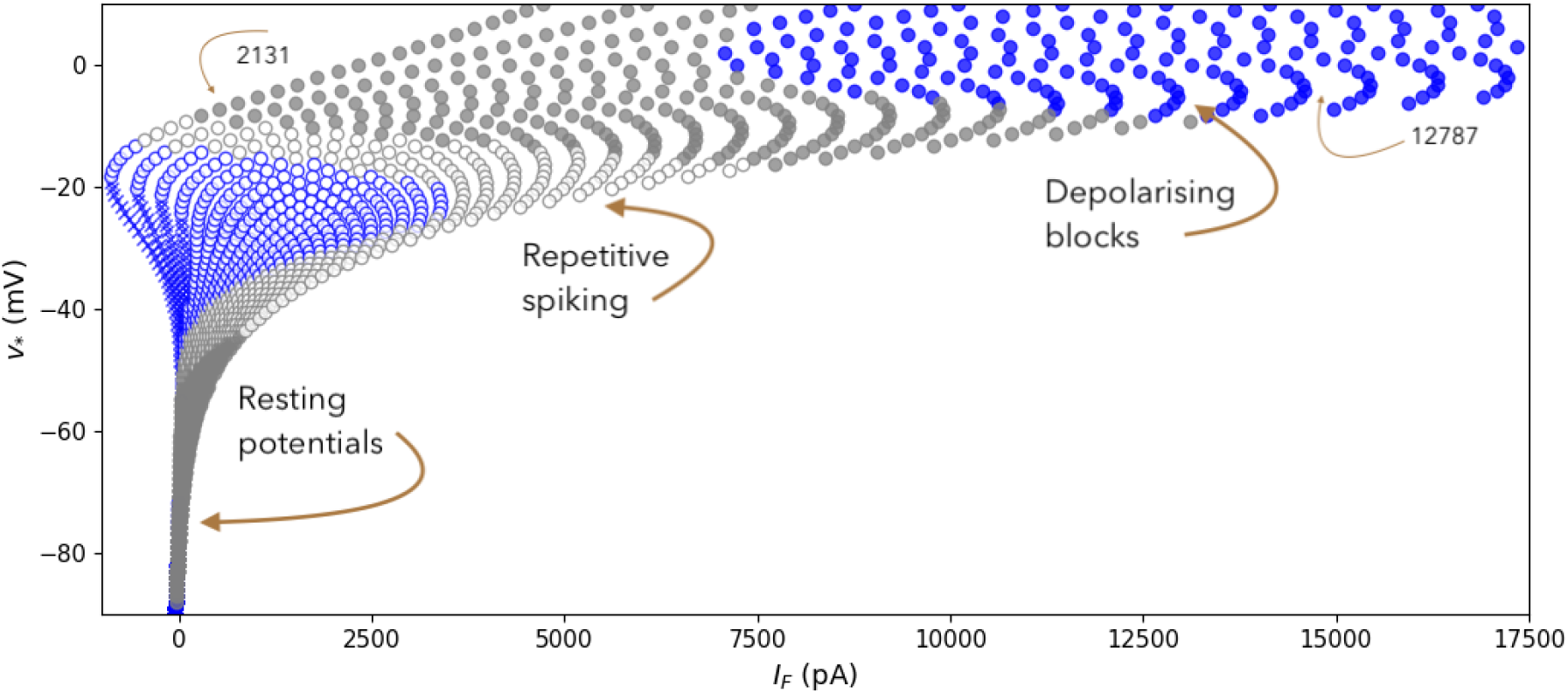
Bifurcation structure of the model for values of *N*_KD_ between 2131 and 10123 while keeping all other parameters constant. Attracting points are represented with colour-filled circles, repelling points with circles filled in white, and saddle nodes as blue crosses. Focus points are shown in grey, nodes are shown in blue. The left-most traces correspond to *N*_KD_=2131. The trace at the far right corresponds to *N*_KD_ = 10123.

#### Integration near resting potential and input currents

Now consider the changes in the fixed points as *I_F_* increases. The fixed points that have the lowest *v_∗_*-values (near *v*_NK_) are attractor nodes at first, and may become attracting foci for larger *I_F_* before becoming unstable (Fig. 4, transition between blue and grey points in any of the fixed point curves). This transition between attracting nodes and attracting foci is effectively a switch between integration modes in which the first mode the convergence to rest is monotonous (aggregation). In the second mode, the convergence to rest is oscillatory, (resonation).

This switch between attracting nodes and attracting foci shows that single neurons are capable of integrating synaptic input in both, aggregating or resonant modes, depending on the size of the input current (Destexhe et al., 2003). Increasing the number of Kv2.1 channels increases the values of *I_F_* for which attractor nodes become foci and the range for *I_F_* in which the attracting foci are present (right-most traces in Fig. 4). That is, increasing the expression of Kv2.1 channels makes neurons more likely to integrate inputs in resonant mode, and increases the range for which input current may cause firing without causing a depolarisation block.

#### Repetitive firing regime

The second major change in the sequence as *I_F_* increases is that the attractivity of the fixed point associated to the resting potential changes to repulsion, and in most cases involving Andronov-Hopf bifurcations that may be supercritical or subcritical. In the last case limit cycles emerge before the attracting fixed point (rest) looses its attractivity and the system becomes bistable with values of *I_F_* within an short interval. The next fixed point bifurcation may be either a switch from repulsive foci to repulsive nodes which may in turn become saddle nodes, or a more direct switch from repulsive foci to saddle nodes (systems with three fixed points). As *N*_KD_ is increased, saddle node points disappear from the curves and the interval for *I_F_* in which it is possible to observe attracting nodes (left-most portion) also decreases in size.

#### Depolarisation block

After repulsive fixed points appear in the fixed point curves, the next bifurcations for larger *I_F_* will include a switch back to attractors (nodes or foci) associated higher values of *v_∗_* near 0. These last fixed point types are those that correspond to depolarisation blocks.

#### 3.2.1 I-clamp dynamics and predictions from bifurcation structures

The qualitative changes in the excitability of autonomous system like the ones presented here can then be observed for different values of one or more of the parameters (e.g. convergence dynamics near different types of attractors). Of particular interest, we consider a 3-phase, box-shaped current clamp experiment with amplitudes *I*_0_, *I*_1_, and then *I*_0_, lasting long enough for the system to be near an attractor by the end of each stimulus phase. The changes in the stimulus amplitude yield 2 dynamical systems *ϕ*_0_ and *ϕ*_1_, respectively (Rasmussen, 2007; Strogatz, 2018), with three initial conditions to consider; two for *ϕ*_0_ and one for *ϕ*_1_. Assume that the dynamical system *ϕ*_0_ starts at an initial condition (*w*_0_*, v*_0_) and it is near the closest attractor of the system just before changing the stimulus current from *I*_0_ to *I*_1_. The trajectory for the (new) dynamical system *ϕ*_1_ after *I*_0_ *→ I*_1_ would then start from an initial condition that is not an attractor in the new system *ϕ*_1_, and move toward the nearest attractor. When the stimulus is set to back to *I*_0_, the system returns to the original *ϕ*_0_ and the trajectory then evolves from near an attractor for *ϕ*_1_, toward its nearest attractor for *ϕ*_0_. If the system *ϕ*_0_ is bistable, the two trajectories for the system *ϕ*_0_ may converge to different attractors depending on whether the initial conditions were located in two different basins of attraction. A behaviour that is often observed experimentally occurs when the first attractor is a fixed point, and the second one is a limit cycle. As a result, *I*_0_ yields convergence to a resting potential in the first interval of the protocol, and repetitive spiking for large enough *I*_1_, and repetitive spiking toward a different limit cycle after the stimulus is set back from *I*_1_ to *I*_0_ (Lee and Heckman, 1998).

Now let us examine the dynamics two model neurons with Kv2.1 channel expression levels resulting in three and one fixed points for *I_F_* = 0, respectively. The idea is to study the transitions between different dynamical regimes for the same neuron, as those induced by increasing the stimulus amplitude during an I-clamp experiment (Fig. 4 and Fig. 5). To interpret our results, we assume that there are no changes in the biological properties of a neuron during the time that an I-clamp experiment is done. That is, we assume no changes in the biophysical properties (parameters) of a model neuron during an I-clamp experiment. As mentioned before, the bifurcation diagrams show that the rheobase increases with larger *N*_KD_ (Fig. 4). This means that it takes more input current to obtain repetitive firing when *N*_KD_ increases, but also, the way in which repetitive firing is generated is more likely to occur in a resonant way because fixed points associated to resting potentials are more likely to be foci if *N*_KD_ (see also Fig. 5A2 and B2).

**Figure 5:**
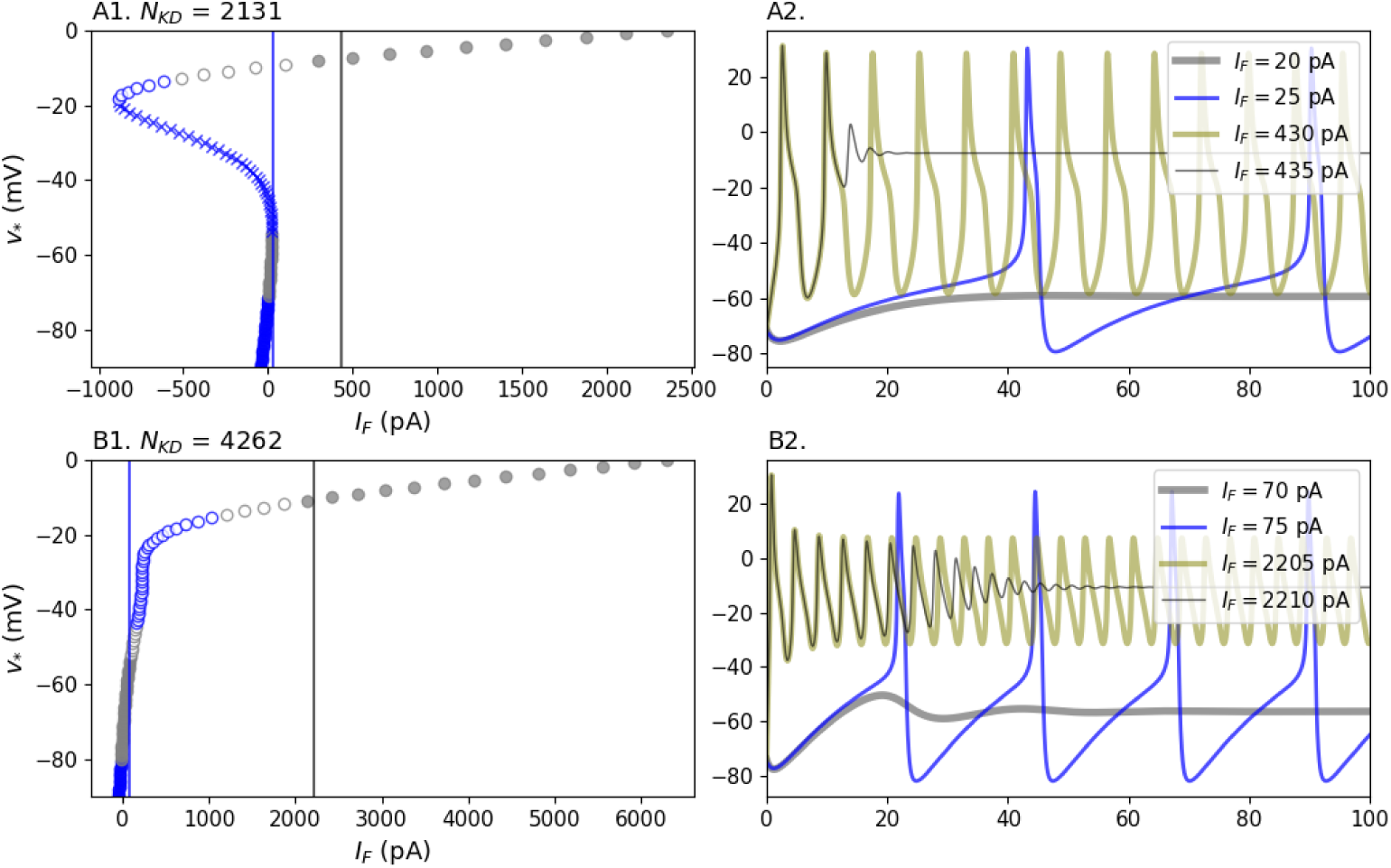
Bifurcations and dynamics for single cell models. (A1) and (B1) show changes in fixed point types occur with *I_F_* (pA) as a bifurcation parameter (see Fig. 2 for comparison). All parameters in the model are identical except for *N*_KD_ *∈* [2131, 4262]. Panels (A2) and (B2) show four representative responses to current clamp stimulation for values of *I_F_* before (grey) and after repetitive spiking (blue), and before and after depolarisation block (olive and black respectively). All trajectories shown for each neuron were obtained from the same initial conditions near rest.

Limit cycles typically exist within and around the region where fixed points are not attractors (e.g. near saddles, or when the *w*-nullcline intersects the *u*-nullcline at its middle branch). However, it is worth noticing that bifurcations involving limit cycles are difficult to detect automatically. Also limit cycle bifurcations are likely to show up in I-clamp protocols, as large enough currents may involve the sudden appearance of limit cycles without changes in the number or types of fixed points (bi-stability, as discussed above).

If *N*_KD_ increases, the range for *I_F_* in which attractor limit cycles occur is larger, and also involves larger input currents (Fig. 5A2 and B2, and Fig. 6). The theoretical lower bound for such a range is the rheobase (Fig. 6). The range for which input current yields repetitive spiking can be loosely identified by finding what values of *I_F_* yield repulsive fixed points within a single bifurcation curve. From there, the *I_F_*-interval for fixed point repulsivity shifts to the right and becomes larger for larger values of *N*_KD_ (see right-most traces in Fig. 4, and compare panels in Fig. 5 and Fig. 6A). In contrast, it is worth noticing that the interval for *v_∗_*-values associated to repetitive spiking (limit cycles) shortens as the number *N*_KD_ increases (Fig. 6B).

**Figure 6:**
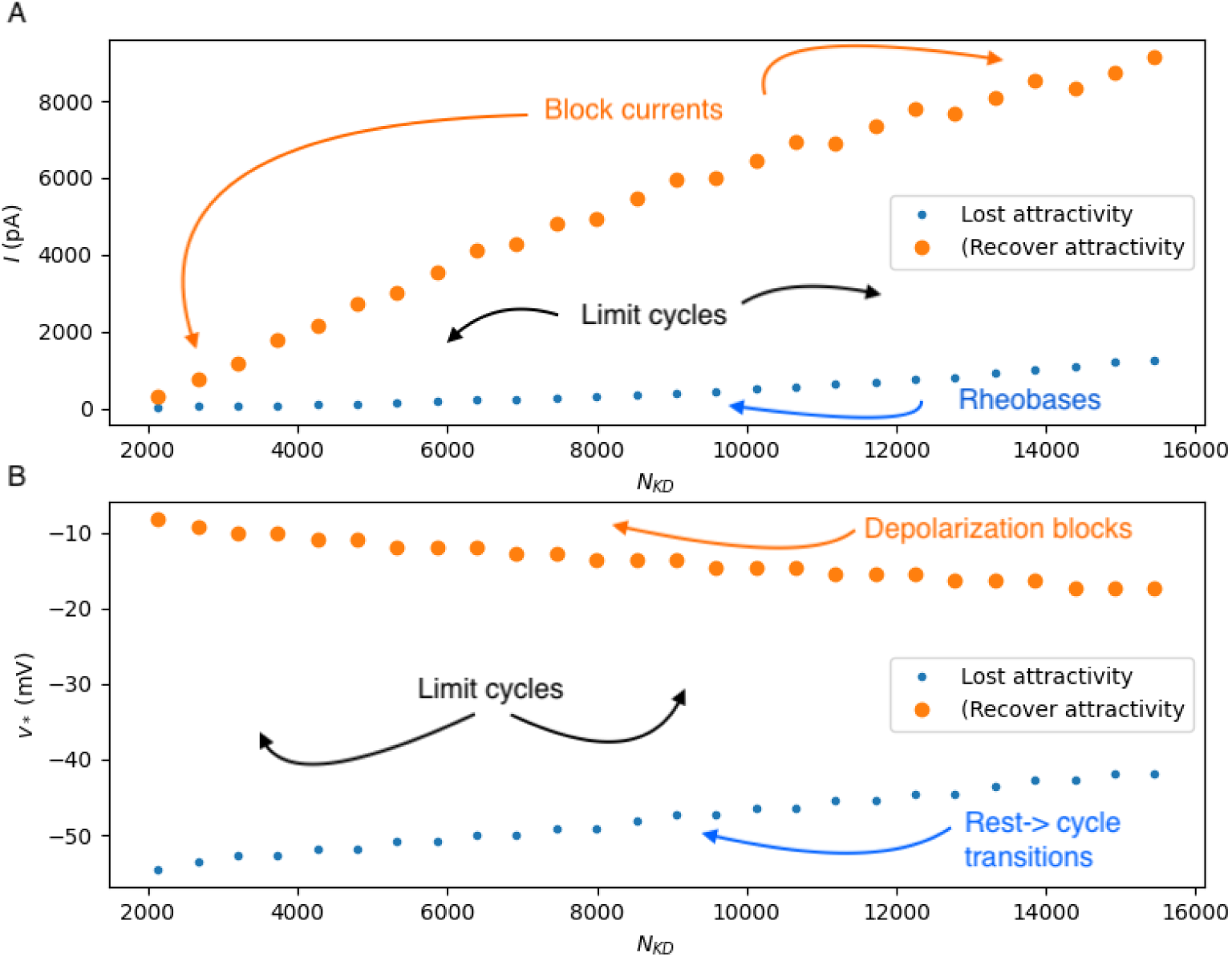
Ranges for the I-Clamp input amplitudes (A) and steady state voltages (B) corresponding to loss (blue dots) and gain (orange dots) of attractivity of the fixed points, respectively (compare with Fig. 4). The lower limit in (A) can be thought of as an approximation to the rheobase current. The upper limit in (B) can be thought of as an approximation to the points where depolarisation blocks occur. The regions where limit cycles occur are pointed to with black arrows in both graphs.

## 4 Discussion

In this article we have taken a solid step into the theoretical study of how K-channel expression changes the excitability of neurons. We do so by systematically studying the geometry and bifurcations associated to Kv2.1 channel expression, using 2D biophysical models of neuronal membrane with the simplest complement of transmembrane currents. This approach allows us to establish solid, basic results such as analytical calculation of fixed point types and from there variation of the level of expression of Kv2.1 channels, before extending our models to include other K-currents that may require higher dimensional systems with more parameters.

There have been other studies taking various approaches that aim to shed light on the general question about the influence of K-channels on membrane excitability (Santini and Porter, 2010; Shipston, 2001), including studies that make spatial considerations (Zang and Marder, 2023). For instance, Drion et al. (2015) investigate how changes in the amplitude of A-type K-channels and CaL channels modify the qualitative features of I-F curves. In that work, Drion et al. use a model with A-type K-currents and L-type Ca-currents to study the influence of K-currents on the classification for excitability based on I-F curves originally proposed by Hodgkin and Huxley (1952). Our approach contrasts with approaches in studies like those of Drion et al. and Zang and Marder in that we first use a reduced 2D biophysical model with the minimum, yet, physiologically meaningful complement of ion currents, to study the excitability of a neuron. In our work, only the presence of delayed-rectifier (non-inactivating) K-channels is varied. This allows us to perform systematic and controlled parameter variations, which can be extended further to include variability that may include other types of K-channels. Further, we do so, performing geometrical analysis, and we draw our conclusions on a classification based on sequences of bifurcations and phase plane analysis.

The question that is dealt with in this article brings out the necessity of a definition for excitability that describes the activity of electrically excitable cells properly; the literature is crowded with attempts that characterise the neuronal electrical activity (Peng and Wu, 2007), but falls short in missing some key features from the physiology or the experiments that cause their theories to fall into contradictions. One notable instance is the incorrect idea that excitability involves bifurcations (e.g. changing the input current, thereby changing the evolution of the dynamical system), or that it can be classified with respect to being near a bifurcation (Izhikevich, 2007). We address the question *en passage*, while studying dynamics like those observed in box-shape I-clamp experiments. The idea is simple: to model the behaviour of a neuron in response to a change in an experimental condition (e.g. *I_F_*), we can change parameters representing the condition, and use a *family* of autonomous dynamical systems (not just one) with the same functional form for the core membrane dynamics, to reproduce the responses to the experimental manipulation. Importantly, this can be done within a time interval split into three phases in which the stimulus amplitude is constant, allowing the systems to remain autonomous. We then link the different behaviours that a single (model) neuron may exhibit with a given pattern of channel expression. The idea was to first reproduce observable experimental behaviours in single-neuron instances of our model, and then try to explain the mechanisms underlying the observations with geometrical analysis. While doing so, we give a working definition of a pulse and from there define the concept of excitability using the geometry of the phase plane. To do that we propose (mathematical) conditions for excitability based on how the direction fields of the phase plane are arranged. After observing that there are only four qualitatively different directions, we propose conditions for excitability that involve the existence of an excitability region (called *R*_1_ for the models presented here, Fig. 1) from which a neuron could produce action potentials. The excitability region may change in shape and size with *I_F_*, increasing, then decreasing in size as *I_F_* increases. The important issue is thus why some initial conditions for the same model neuron result in APs and some do not? Once addressed, the non-existence of voltage or current thresholds can be explained as a corollary by focusing on the location of the initial conditions. From the fact that the two nullclines of the system can be regarded as the graphs of the functions *Z_w_* and *Z_u_* of *u* (and therefore of *v* after the appropriate re-scaling), we can conjecture another corollary: assume that *I_F_* is such that the fixed point with the lowest *v*-value occurs in the first branch of the *v*-nullcline (the first branch that is decreasing as a function of *v* for *v > v*_KD_). Then there is a continuous curve that defines the upper boundary of the excitability region in the *w* direction, which can be used to predict whether an initial condition will result in an action potential. The curve is composed of those points (*w, v*) on the nullclines, with *w* taking the minimum positive value (equation (17), Fig. 1, curve indicated by black dots).

This analysis also yields an interesting way of thinking about decreasing the excitability of a neuron. In short, any parameter that decreases the area of the (excitability) region *R*_1_ is a candidate to decrease excitability. In particular, more Kv2.1 channels are expressed in the membrane, the size of the region *R*_1_ within the domain *D* = (0, 1) *×* (*v*_KD_*, v*_NT_) decreases. If we were to select initial conditions randomly, decreasing the size of the *R*_1_ region translates into a smaller probability of producing APs. This suggest a general way to address whether the expression of channels can affect how neurons produce action potentials, thereby linking bifurcation sequences to (biophysical) parameter changes involving channel expression; a impossible task to perform in experiments. In this regard, our main result is that increasing the number of Kv2.1 potassium channels in the neuronal membrane changes the way in which neurons respond to input currents in at least three ways: it makes resonant integration of inputs more likely to occur, it increases the range in which input current causes repetitive firing (Fig. 6A), but it does so at the expense of making it more difficult for the neuron to fire or further, to become continuously excited because it also increases the rheobase. This is relevant for highly active networks, in which increasing the expression of K-channels is one of the neuronal mechanisms that have been observed to regulate input-output responses (Lee et al., 2015).

The overall behaviour of the system for larger values of *N*_KD_ is in agreement with the intuitive idea that it is more difficult to produce APs if the number of K^+^channels in the membrane is larger. However, it is not obvious that the range for input current for which APs occur becomes larger. Our analysis also explains results from an earlier study (Herrera-Valdez et al., 2013) in which recordings from adult *Drosophila melanogaster* MN5 neurons displayed depolarisation blocks for relatively large input currents under I-clamp, in many cases without being able to display transitions into repetitive spiking, or having done so for only large input currents within a short range of amplitudes. Our analysis gives rise to a simple hypothesis that could be tested experimentally: the number of Shab channels in those neurons that had the tendency of becoming block-depolarised was very large. This would be in line with the idea that the recruitment of MN5 and its contractile command over the dorsolateral wing muscle occurs under very tight control.

### 4.1 On the necessity of improving our biophysical models of membrane potential

*Mathematical modelling* has been profusely used to study transmembrane transport and the electrical signalling that results from it.

To date, the largest body of work in the mathematical modelling of for the membrane potential in excitable cells derived from an equivalent circuit in the seminal work of Hodgkin and Huxley (1952), and the studies by other researchers that followed. In short, the time-dependent change in membrane potential *v* is given by *C_m_∂_t_* = *I_F_ −*∑*_x_ I_x_* where *C_m_* represents a “membrane capacitance”, and *I_x_* represents a generic transmembrane current given by 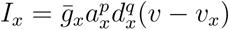, where the difference (*v − v_x_*) can be thought of as an “Ohmic” driving force for voltage. The term ḡ*_x_* represents a maximal “conductance” modulated by variables *a_x_* and *b_x_* between 0 and 1 that represent proportions of “activated” and “de-inactivated” channels (the latter confusedly called “inactivation” variable). The voltage-dependent dynamics of *a_x_* and *b_x_* are linear with coefficients that depend on *v*, and the powers *p* and *q* allow to statistically fit the non-linear components of gating as analysed in the experiments by Hodgkin and Huxley and others that followed the same logic and interpretation (see Aldrich et al. (1983)).

However, there are, at least three issues of critical conceptual importance from the original formulation by Hodgkin and Huxley (1952) that should no longer be ignored based on current knowledge. First, the change in membrane potential is based on an “equivalent circuit” formulation in which the conducting media around the membrane and the transmembrane proteins are respectively regarded as a capacitor and resistors arranged in parallel. In principle, cell membranes can be thought of as non-conductive space between conductive media similar to the air separating metal plates in the classical capacitor. However, there is evidence that ions (and their charge) accumulate non-linearly around the membrane (Everitt and Haydon, 1968; Wobschall, 1972), in contrast with the linear charge accumulation in classical electrical capacitors (Giancoli, 2000). Nevertheless, it is possible to derive an equation describing the change in transmembrane potential from basic biophysical principles without involving equivalent circuits (Herrera-Valdez, 2020). The derivation shows *en passage* that the “capacitance” from the equivalent circuit formulation can be obtained by assuming that the voltage-dependence of charge density around the membrane is linear. Also, there is a generic formulation to model transmembrane transport mediated by both channels and pumps that is based on thermodynamical principles (Herrera-Valdez, 2018). In that work, there is a simple mathematical demonstration of how a Taylor approximation around the reversal potential in the general formula for current yields the “conductance” from the Hodgkin and Huxley model after truncating to first order. Second, transmembrane currents carried through channels are electrodiffusive, not Ohmic. The “conductance” can be shown to emerge from calculating a first-order Taylor approximation of expressions that do describe electrodiffusion (Herrera-Valdez, 2018). In other words, the conductance-based approach is only a linear approximation around the reversal potential for a current, and it disregards first principles descriptions of electrodiffusive fluxes of ions (e.g. Nernst-Planck). Further, the voltage-current relationships reconstructed from single channel recordings are clearly not linear, as predicted by the conductance-based approach (Bittner and Hanck, 2008; Vandenberg and Bezanilla, 1991).

Also, the powers *p* and *q* in the gating terms 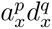 of the Hodgkin and Huxley (1952) model reduce the size of the total current amplitude and the (whole-cell) reduction is automatically compensated during the fit by choosing larger values for *ḡ_x_*. This is important because *ḡ_x_* can be thought of as *N_x_g_x_* where *N_x_* represents the number of *x*-channels and *g_x_* the “conductance” calculated from single-channel recordings. With this description, the Hodgkin and Huxley model predicts that the number of Na^+^channels is approximately 3 times as large as the number of K^+^channels in the membrane, which is opposite to evidence obtained recently with more advanced techniques (Schulz et al., 2007). A fourth issue that emerged over the years in an attempt to overcome the shortcomings of the conductance-based formulation is that there is a very large number of different descriptions (functional forms) for transmembrane transport mechanisms mediated by channels and pumps. Examples can be observed in modelling studies by DiFrancesco and Noble (1985) and Rasmusson et al. (1990), to name some. There is also a more appropriate and logical way of modelling the evolution rule for channel gating at the (channel) population level: a function with logistic form, which naturally yields solutions like those obtained from voltage-clamp experiments (Herrera-Valdez, 2012b). The logistic-like evolution is more suitable for populations of channels undergoing similar conformational changes, and avoids the use of powers in gating terms with the subsequent decrease in the parameters describing the numbers of channels in the membrane.

### 4.2 Future directions and conclusions

One direction of possible future work involves relaxing the assumption in our model that all transmembrane concentrations of ions and the ATP/ADP ratio are constant (see Johnston et al., 1995, chapter 2, example 2.1). The relevance of this assumption may be tested, as our model is intended to apply to small and large compartments alike, but the transmembrane ionic concentrations may change drastically in small periods of time in small compartments. It would be important to extend our work in that direction following the steps of Barreto and Cressman among other researchers. In particular, it could be interesting to compare the results presented here with those obtained using phenomenological models of Na-K ATPase activity (Hasenstaub et al., 2010; Le Masson et al., 2014; Zang and De Schutter, 2021).

One advantage of the newer modelling schemes used here is that the formulations are biophysical by construction, without unwarranted statistical fits, or other construction issues that could result in difficulties interpreting results consistently with respect to biological facts. In addition to the biophysical nature of our models, we have also been able to perform some tasks analytically, including the calculation of the eigenvalues of the Jacobian matrix evaluated at fixed points, their real and imaginary parts, and the subsequent fixed point types. Another future direction of our work will include exploring mathematical aspects of the theory that involve explicit calculations and linking those calculations to the physiology of excitability. Another slightly different direction of particular interest is to use models like the one presented here to systematically link the transcription of genes coding for different proteins, and the effect this may have on the physiological function of systems of interest.

## Acknowledgements

The work of MAHV was supported by DGAPA-UNAM grant IN-228820.

## Footnotes

1 *k* represents Boltzmann constant (1.380649 *·* 10^−23^ J/K), *T* the absolute temperature (K), and *q* is the elementary charge (1.602176634*·* 10^−19^ Coulombs)

## References

David J. Aidley. The Physiology of Excitable Cells. Cambridge University Press, 4 edition, 1998. ISBN 0521574153, 9780521574150. URL http://gen.lib.rus.ec/book/index.php?md5=25AD083C33F37AC8E44F3CE90E3B3B84.

R. W. Aldrich, D. P. Corey, and C. F. Stevens. A reinterpretation of mammalian sodium channel gating based on single channel recording. Nature, 306(5942):436–441, 1983.

Frances M Ashcroft. Atp-sensitive potassium channelopathies: focus on insulin secretion. The Journal of clinical investigation, 115(8):2047–2058, 2005.

E. Av-Ron. The role of a transient potassium current in a bursting neuron model. Journal of Mathematical Biology, 33(1):71–87, 1994.

E. Av-Ron, H. Parnas, and L. A. Segel. A minimal biophysical model for an excitable and oscillatory neuron. Biological Cybernetics, 65(6):487–500, 1991.

Evyatar Av-Ron, Hanna Parnas, and Lee A. Segel. A basic biophysical model for bursting neurons. Biological Cybernetics, 69:87–95, 1993.

Ernest Barreto and John R Cressman. Ion concentration dynamics as a mechanism for neuronal bursting. Journal of biological physics, 37:361–373, 2011.

B. P. Bean and E. Rios. Nonlinear charge movement in mammalian cardiac ventricular cells. components from na and ca channel gating. The Journal of general physiology, 94(1):65, 1989.

Bruce P Bean. The action potential in mammalian central neurons. Nature Reviews Neuroscience, 8(6):451–465, 2007.

Eduardo E Benarroch. Na+, k+-atpase: functions in the nervous system and involvement in neurologic disease. Neurology, 76(3):287–293, 2011.

Francisco Bezanilla. Voltage-gated ion channels. Biological Membrane Ion Channels: Dynamics, Structure, and Applications, pages 81–118, 2007.

Katie C Bittner and Dorothy A Hanck. The relationship between single-channel and whole-cell conductance in the t-type ca2+ channel cav3.1. Biophysical journal, 95(2):931–941, 2008.

M.P. Blaustein, J.P.Y. Kao, and D.R. Matteson. Cellular physiology. Elsevier/Mosby, 2004. ISBN 0323013414.

Brett C Carter and Bruce P Bean. Sodium entry during action potentials of mammalian neurons: incomplete inactivation and reduced metabolic efficiency in fast-spiking neurons. Neuron, 64(6): 898–909, 2009.

J. B. Chapman. The reversal potential for an electrogenic sodium pump. a method for determining the free energy of atp breakdown? 72:403–408, 1978.

J Brian Chapman. On the reversibility of the sodium pump in dialyzed squid axons: A method for determining the free energy of atp breakdown? The Journal of general physiology, 62(5):643, 1973.

Senyon Choe. Potassium channel structures. Nature Reviews Neuroscience, 3(2):115–121, 2002.

Graham L Collingridge, Richard W Olsen, John Peters, and Michael Spedding. A nomenclature for ligand-gated ion channels. Neuropharmacology, 56(1):2–5, 2009.

M Covarrubias, A Wei, and L Salkoff. Shaker, Shal, Shab, and Shaw expresss independent K-current systems. Neuron(Cambridge, Mass.), 7(5):763–773, 1991.

Alain Destexhe, Michael Rudolph, and Denis Paré. The high-conductance state of neocortical neurons in vivo. Nature reviews neuroscience, 4(9):739–751, 2003.

D. DiFrancesco and D. Noble. A model of cardiac electrical activity incorporating ionic pumps and concentration changes. *Philosophical Transactions of the Royal Society of London. Series B*, Biological Sciences, 307:353–398, 1985.

Guillaume Drion, Timothy O’Leary, and Eve Marder. Ion channel degeneracy enables robust and tunable neuronal firing rates. Proceedings of the National Academy of Sciences, 112(38): E5361–E5370, 2015.

S. E. Dryer. Na+-activated K+ channels: a new family of large-conductance ion channels. Trends in neurosciences, 17(4):155–160, 1994.

G. B. Ermentrout and D. Terman. Foundations of mathematical neuroscience (Interdisciplinary applied mathematics series, Vol. 35). Springer Verlag, 2010.

CT Everitt and DA Haydon. Electrical capacitance of a lipid membrane separating two aqueous phases. Journal of theoretical biology, 18(3):371–379, 1968.

R Douglas Fields. Oligodendrocytes changing the rules: action potentials in glia and oligodendrocytes controlling action potentials. The Neuroscientist, 14(6):540–543, 2008.

Richard FitzHugh. Impulses and physiological states in theoretical models of nerve membrane. Biophysical journal, 1(6):445–466, 1961.

D.C. Gadsby. Ion channels versus ion pumps: the principal difference, in principle. Nature Reviews Molecular Cell Biology, 10(5):344–352, 2009.

Douglas C Giancoli. Physics for scientists and engineers third edition, 2000.

R Granit, D Kernell, and Shortess G. K. Quantitative aspects of repetitive firing of mammalian motoneurones, caused by injected currents. Journal of Physiology, 168:911–931, 1963a.

R. Granit, D. Kernell, and R. S. Smith. Delayed depolarization and the repetitive response to intracellular stimulation of mammalian motoneurones. The Journal of Physiology, 168(4):890, 1963b.

Andrea Hasenstaub, Stephani Otte, Edward Callaway, and Terrence J Sejnowski. Metabolic cost as a unifying principle governing neuronal biophysics. Proceedings of the National Academy of Sciences, 107(27):12329–12334, 2010.

Marco Arieli Herrera-Valdez. Membranes with the same ion channel populations but different excitabilities. PloS one, 7(4):e34636, 2012a.

Marco Arieli Herrera-Valdez. Same ion channel populations and different excitabilities: beyond the conductance-based mod el. BMC Neuroscience, 13(Suppl 1):P120, 2012b.

Marco Arieli Herrera-Valdez. Geometry and nonlinear dynamics underlying electrophysiological phenotypes in biophysical models of membrane potential. Dissertation., 2014.

Marco Arieli Herrera-Valdez. A thermodynamic description for physiological transmembrane transport [version 2; referees: 2 approved]. F1000Research, 7(1468), 2018. doi: 10.12688/f1000research.16169.2.

Marco Arieli Herrera-Valdez. An equation for the biological transmembrane potential from basic biophysical principles. F1000Research, 9(676):676, 2020.

Marco Arieli Herrera-Valdez and Joceline Lega. Reduced models for the pacemaker dynamics of cardiac cells. Journal of Theoretical Biology, 270(1):164–176, 2011.

Marco Arieli Herrera-Valdez, Erin Christy McKiernan, Sandra Daniela Berger, Stephanie Ryglewski, Carsten Duch, and Sharon Crook. Relating ion channel expression, bifurcation structure, and diverse firing patterns in a model of an identified motor neuron. Journal of Computational Neuroscience, pages 1–19, 2013.

Stephan Heyse, Ingo Wuddel, Hans-Jürgen Apell, and Werner Stürmer. Partial reactions of the na, k-atpase: determination of rate constants. The Journal of general physiology, 104(2):197–240, 1994.

B Hille. *Ionic Channels of Excitable Membranes*. Sinauer Associates, Sinauer Associates, Inc. Sunderland, Mass. 01375, 1992.

A. L. Hodgkin and A. F. Huxley. A quantitative description of membrane current and its application to conduction and excitation in nerve. Journal of Physiology, 117:500–544, 1952.

At L Hodgkin and Pv Horowicz. The influence of potassium and chloride ions on the membrane potential of single muscle fibres. The Journal of physiology, 148(1):127, 1959.

J. Hounsgaard, H. Hultborn, B. Jespersen, and O. Kiehn. Intrinsic membrane properties causing a bistable behaviour of *α*-motoneurones. Experimental brain research, 55(2):391–394, 1984.

Eugene Izhikevich. Dynamical Systems in Neuroscience. MIT, MIT Press, 55 Hayward Street, Cambridge MA 02142, 2007.

D. Johnston, S. M. S. Wu, and R. Gray. Foundations of cellular neurophysiology. MIT press Cambridge, MA, 1995. ISBN 0262100533.

Siyavash Joukar. A comparative review on heart ion channels, action potentials and electrocardiogram in rodents and human: extrapolation of experimental insights to clinic. Laboratory Animal Research, 37(1):1–15, 2021.

Carsten Juel. Muscle action potential propagation velocity changes during activity. Muscle & Nerve: Official Journal of the American Association of Electrodiagnostic Medicine, 11(7):714–719, 1988.

Gwendal Le Masson, Serge Przedborski, and LF4167823 Abbott. A computational model of motor neuron degeneration. Neuron, 83(4):975–988, 2014.

Kwan Young Lee, Sara E Royston, Max O Vest, Daniel J Ley, Seungbae Lee, Eric C Bolton, and Hee Jung Chung. N-methyl-d-aspartate receptors mediate activity-dependent down-regulation of potassium channel genes during the expression of homeostatic intrinsic plasticity. Molecular brain, 8(1):1–16, 2015.

R. H. Lee and C. J. Heckman. Bistability in spinal motoneurons in vivo: systematic variations in persistent inward currents. Journal of neurophysiology, 80(2):583, 1998.

W. H. Lin, D. E. Wright, N. I. Muraro, and R. A. Baines. Alternative Splicing in the Voltage-Gated Sodium Channel DmNav Regulates Activation, Inactivation, and Persistent Current. Journal of Neurophysiology, 102(3):1994, 2009.

MJ Mason, AK Simpson, MP Mahaut-Smith, and HPC Robinson. The interpretation of current-clamp recordings in the cell-attached patch-clamp configuration. Biophysical journal, 88(1): 739–750, 2005.

E. K. Mattews and Y. Sakamoto. Pancreatic islet cells: electrogenic and electrodiffusional control of membrane potential. The Journal of Physiology, 246(2):439, 1975. ISSN 0022-3751.

Hiroaki Misonou, Durga P Mohapatra, and James S Trimmer. Kv2. 1: a voltage-gated k+ channel critical to dynamic control of neuronal excitability. Neurotoxicology, 26(5):743–752, 2005.

Durga P Mohapatra, Hiroaki Misonou, Pan Sheng-Jun, Joshua E Held, D James Surmeier, and James S Trimmer. Regulation of intrinsic excitability in hippocampal neurons by activity-dependent modulation of the kv2. 1 potassium channel. Channels, 3(1):46–56, 2009.

Mitsutoshi Munakata, Mika Fujimoto, Young-Ho Jin, and Norio Akaike. Characterization of electrogenic na/k pump in rat neostriatal neurons. Brain research, 800(2):282–293, 1998.

Hideyuki Murakoshi and James S Trimmer. Identification of the kv2. 1 k+ channel as a major component of the delayed rectifier k+ current in rat hippocampal neurons. Journal of Neuroscience, 19(5):1728–1735, 1999.

Shigehiro Nakajima, Shizuko Iwasaki, and Kunihiko Obata. Delayed rectification and anomalous rectification in skeletal muscle membrane. Proceedings of the Japan Academy, 37(8):505–508, 1961.

Shigehiro Nakajima, Shizuko Iwasaki, and Kunihiko Obata. Delayed rectification and anomalous rectification in frog’s skeletal muscle membrane. The Journal of general physiology, 46(1): 97–115, 1962.

Björn Naundorf, Fred Wolf, and Maxim Volgushev. Unique features of action potential initiation in cortical neurons. Nature, 440(7087):1060, 2006.

I-Feng Peng and Chun-Fang Wu. Differential contributions of Shaker and Shab K+ currents to neuronal firing patterns in Drosophila. Journal of Neurophysiology, 97(1):780, 2007.

Ole H Petersen and Yoshio Maruyama. Calcium-activated potassium channels and their role in secretion. Nature, 307(5953):693–696, 1984.

Martin Rasmussen. Attractivity and bifurcation for nonautonomous dynamical systems. Springer, 2007.

R. L. Rasmusson, J. W. Clark, W. R. Giles, E. F. Shibata, and D. L. Campbell. A mathematical model of bullfrog cardiac pacemaker cell. Am. J. Physiol., 259:H352–H369, 1990.

J. Rinzel. Excitation dynamics: insights from simplified membrane models. In Fed. Proc. 44, volume 2944, 1985.

J. Rinzel and G. B. Ermentrout. Analysis of neural excitability and oscillations, Methods in neuronal modeling: From synapses to networks, 1989.

John Rinzel and G Bard Ermentrout. Analysis of neural excitability and oscillations. Methods in neuronal modeling, 2:251–292, 1998.

Pankaj Sah. Ca2+-activated k+ currents in neurones: types, physiological roles and modulation. Trends in neurosciences, 19(4):150–154, 1996.

L. Salkoff, K. Baker, A. Butler, M. Covarrubias, M. D. Pak, and A. Wei. An essential set of K+ channels conserved in flies, mice and humans. Trends in Neurosciences, 15(5):161–166, 1992.

Edwin Santini and James T Porter. M-type potassium channels modulate the intrinsic excitability of infralimbic neurons and regulate fear expression and extinction. Journal of Neuroscience, 30 (37):12379–12386, 2010.

David J Schulz, Jean-Marc Goaillard, and Eve E Marder. Quantitative expression profiling of identified neurons reveals cell-specific constraints on highly variable levels of gene expression. Proceedings of the National Academy of Sciences, 104(32):13187–13191, 2007.

Liu Shi, Chaoli Mu, Tao Gao, Wenxin Chai, Anzhi Sheng, Tianshu Chen, Jie Yang, Xiaoli Zhu, and Genxi Li. Rhodopsin-like ionic gate fabricated with graphene oxide and isomeric dna switch for efficient photocontrol of ion transport. Journal of the American Chemical Society, 141(20): 8239–8243, 2019.

M.J. Shipston. Alternative splicing of potassium channels: a dynamic switch of cellular excitability. Trends in cell biology, 11(9):353–358, 2001.

Takao Sibaoka. Rapid plant movements triggered by action potentials. The botanical magazine= Shokubutsu-gaku-zasshi, 104:73–95, 1991.

Jens Christian Skou and Mikael Esmann. The na, k-atpase. Journal of bioenergetics and biomembranes, 24:249–261, 1992.

David J Speca, Genki Ogata, Danielle Mandikian, Hannah I Bishop, Steve W Wiler, KJWenzelH Eum, H Jürgen Wenzel, Emily T Doisy, Lucas Matt, Katharine L Campi, et al. Deletion of the kv2. 1 delayed rectifier potassium channel leads to neuronal and behavioral hyperexcitability. Genes, Brain and Behavior, 13(4):394–408, 2014.

Wilfred D Stein and Thomas Litman. Channels, carriers, and pumps: an introduction to membrane transport. Elsevier, 2014.

Steven H Strogatz. Nonlinear dynamics and chaos with student solutions manual: With applications to physics, biology, chemistry, and engineering. CRC press, 2018.

Greg Stuart and Michael Häusser. Initiation and spread of sodium action potentials in cerebellar purkinje cells. Neuron, 13(3):703–712, 1994.

Gürol M Süel, Jordi Garcia-Ojalvo, Louisa M Liberman, and Michael B Elowitz. An excitable gene regulatory circuit induces transient cellular differentiation. Nature, 440(7083):545–550, 2006.

Gurol M Suel, Rajan P Kulkarni, Jonathan Dworkin, Jordi Garcia-Ojalvo, and Michael B Elowitz. Tunability and noise dependence in differentiation dynamics. Science, 315(5819):1716–1719, 2007.

Benjamin A Suter, Michele Migliore, and Gordon MG Shepherd. Intrinsic electrophysiology of mouse corticospinal neurons: a class-specific triad of spike-related properties. Cerebral cortex, 23(8):1965–1977, 2013.

S. Tsunoda and L. Salkoff. The major delayed rectifier in both Drosophila neurons and muscle is encoded by Shab. Journal of Neuroscience, 15(7):5209–5221, 1995.

C. A. Vandenberg and F. Bezanilla. Single-channel, macroscopic, and gating currents from sodium channels in the squid giant axon. Biophysical journal, 60(6):1499–1510, 1991.

Valerie Villiere and Elspeth M McLachlan. Electrophysiological properties of neurons in intact rat dorsal root ganglia classified by conduction velocity and action potential duration. Journal of Neurophysiology, 76(3):1924–1941, 1996.

Yi-Chi Wang, Jyh-Jeen Yang, and Rong-Chi Huang. Intracellular na+ and metabolic modulation of na/k pump and excitability in the rat suprachiasmatic nucleus neurons. Journal of Neurophysiology, 108(7):2024–2032, 2012.

Z Galen Wo and Robert E Oswald. Unraveling the modular design of glutamate-gated ion channels. Trends in neurosciences, 18(4):161–168, 1995.

Darold Wobschall. Voltage dependence of bilayer membrane capacitance. Journal of Colloid and Interface Science, 40(3):417–423, 1972.

Stephen H Wright. Generation of resting membrane potential. Advances in physiology education, 28(4):139–142, 2004.

Yunliang Zang and Erik De Schutter. The cellular electrophysiological properties underlying multiplexed coding in purkinje cells. Journal of Neuroscience, 41(9):1850–1863, 2021.

Yunliang Zang and Eve Marder. Neuronal morphology enhances robustness to perturbations of channel densities. Proceedings of the National Academy of Sciences, 120(8):e2219049120, 2023.

